# Geroprotective drug discovery with an AI-enabled assay for heat resistance

**DOI:** 10.64898/2026.08.10.743574

**Authors:** Thomas J. Hodder, Allancer D.C. Nunes, Bryan A. Martinez, Karla J. Opperman, Paul D. Robbins, Chad L. Myers, Matthew S. Gill

**Affiliations:** Masonic Institute on the Biology of Aging & Metabolism, University of Minnesota; Department of Cell Biology, Genetics & Development, University of Minnesota; Department of Biochemistry, Molecular Biology & Biophysics, University of Minnesota; Department of Computer Science & Engineering, University of Minnesota

**Author notes:** Correspondence: Matthew S. Gill, Chad L. Myers.

## Abstract

Drugs that slow the rate of organismal aging (geroprotectors) have the potential to improve human healthspan by preventing the development of multiple chronic diseases. The nematode *C. elegans* is a proven system for identifying anti-aging drugs, but methods for candidate nomination that scale to high throughput remain limited and have not been widely adopted. To accelerate the discovery process, we have developed an open-source AI-enabled screening platform that provides a rapid, automated, posture-based score of *C. elegans* survival. We used this workflow to screen a library of 2,782 FDA-approved drugs and identified 31 compounds that reproducibly increase heat stress resistance, a known predictor of longevity. Follow-up studies confirmed that many of these compounds confer lifespan extension in worms, and several compounds also have anti-senescent activity in human cells. This simplified and adaptable AI-enabled workflow therefore has the potential to accelerate the discovery of translatable drugs that slow aging.

## Introduction

Since the discovery of the first single-gene mutations that confer extended lifespan in C. elegans (1, 2) numerous other genetic (3, 4) and chemical (5–8) interventions have been discovered that promote longevity. These studies have not only established molecular mechanisms that are causally associated with aging, but they have demonstrated that aging is plastic and amenable to manipulation. The geroscience hypothesis (9) proposes that therapeutic targeting of aging will broadly prevent age-related disease and disorders such as cancer, diabetes, and heart failure. Thus, drugs that target basic mechanisms of aging, termed “geroprotectors”, may serve as a strategy to optimize human healthspan and reduce the societal burden of age-related disease.

It is not clear whether the traditional single-validated-target approach to drug identification is optimal for promoting longevity, as aging is impacted by multiple pathways and the identification of the key target in each is not trivial. An alternative is to focus on the aging phenotype and remain agnostic to the molecular targets. In this respect, high-throughput phenotypic drug screens for longevity in worms have led to the identification of novel compounds that may protect against aging (*5*, *7*, *8*), demonstrating the utility of *C. elegans* as a model system for drug discovery in aging.

However, there remains a need to discover more candidates given the likely rate of attrition in translating drugs from the nematode to mammalian models. One disadvantage of lifespan screens in worms is that survival assays are time-consuming, and thus, we sought to reduce the assay length from several weeks to a few days. To this end, we focused on developing a screening platform for measuring heat stress resistance, which has a proven track record as a predictor of extended lifespan. Following the discovery that long-lived mutant animals are highly thermotolerant (*10*), varied interventions that increase heat stress resistance (*11*, *6*) have shown an association with longevity.

The standard approach to scoring survival in *C. elegans* remains the manual assessment of touch-provoked movement. This creates a major bottleneck in scaling survival assays to a multi-well plate format, and an ideal assay readout would be fully automated to reduce labor and decrease technical variation. Automated readouts of survival have been developed including camera-based movement (*12*), flatbed scanners (*13*), and fluorescent reporter readouts using SYTOX dye (*14*, *15*). However, since many of these approaches require specialized equipment and expertise, we were motivated to develop a simple automated score. Live and dead worms tend to be visually distinct, with a sinusoidal posture in live worms and a straightened posture in dead worms. We hypothesized that these postures could be objectively quantified with a neural network-based classifier (*16*), removing the need for dyes or video analysis for survival quantification and hit detection.

AI-based computer vision workflows have exploded in popularity since the design of AlexNet in 2012 (*17*). Although several deep neural network based workflows for analysis of worms have been developed since then (*18*, *19*), application of these methods to high-throughput drug screens have been limited. Reasons for this may include inflexibility of software outside of a specific problem and difficulty of data collection due to hardware/laboratory requirements for customized equipment. To create an accessible and scalable workflow, we reasoned that the design goal should be to simplify the laboratory protocols and software usage. Rather than developing a customized hardware setup for imaging, we opted to use a standard stage-motorized microscope for image acquisition since this instrumentation is available at most academic and industrial research institutions. The ability to score survival from brightfield images offers multiple advantages over a fluorescent dye-based workflow. In particular, interpretable analysis of fluorescence images in high-content screens typically requires a wash step (*15*) since debris (worm progeny/eggs, bacteria, precipitated drug) can interfere with the readout.

Using an integrated AI-based workflow that quantifies worm posture and heat stress resistance as an assay readout, we have reduced the length of a round of drug screening from multiple weeks to a two-day heat shock and recovery. To demonstrate the utility of this approach we have screened a library of 2,782 drugs for heat stress resistance, and identified 18 drugs that significantly extend lifespan, of which 14 have not previously been implicated in longevity.

## Results

### Development of a scalable, automated workflow for scoring survival

The image-scoring workflow can be separated into three steps: 1) detection, 2) segmentation, and 3) classification. Worms are detected with Faster R-CNN (*20*) with a ResNet18 (*16*) backbone, finetuned on bounding box-annotated images. Bounding-box outputs of the detection step are then used as prompts for Segment Anything Model 2 (SAM 2) **(Fig. 1A)** (*21*) which predicts a mask of the focal object of each bounding box. Masks are made a single color so that the model is forced to use only worm shape in the classification. Comparison of the number of automatically detected worms to manual counts across a range of population sizes demonstrates accurate worm detection by Faster R-CNN **(Fig. 1B)**.

**Figure 1.**
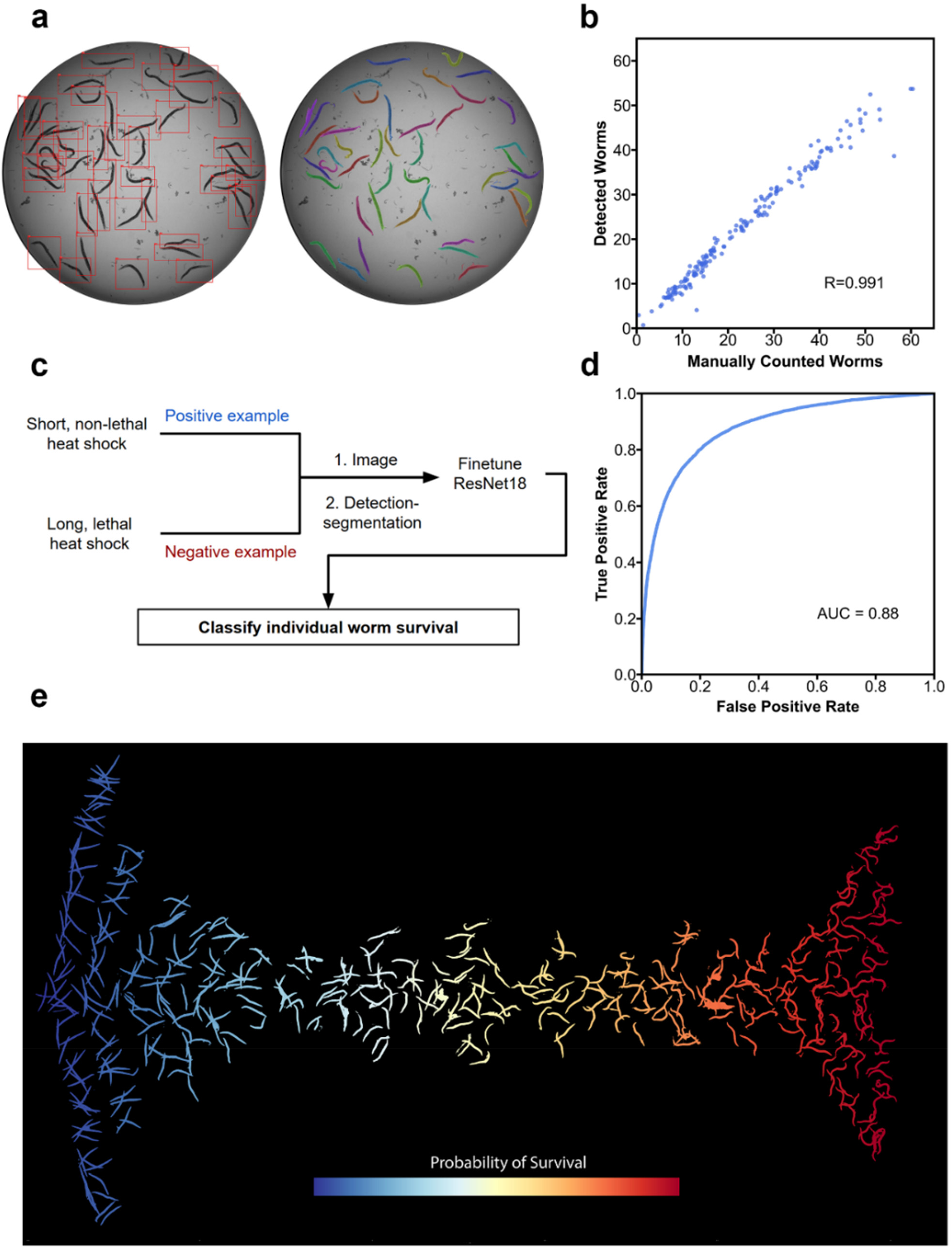
Building a framework for automated worm scoring. (a) Visual demonstration of detection and segmentation. Different colors represent unique worm predictions. (b) Number of worms manually counted by observation of well images versus number of worms detected with a finetuned Faster R-CNN model from the same image. (c) Training and cross validation overview. Three biological replicates of a heat shock with multiple timepoints were performed. For each fold, one biological replicate was held out and the model was trained on masks from the remaining two replicates. (d) Receiver-operator characteristic curve constructed by concatenation of labels and predictions from all three folds. (e) Randomly sampled worm masks from a time course experiment colored by probability of survival assigned by the classifier.

We then fine-tuned a ResNet18 backbone for classification of worm masks to predict the probability of survival of each detected worm. Worms were subjected to heat shocks at various timepoints to derive positive (nonlethal heat shock exposure) and negative (severe heat shock exposure) examples of worms and then imaged for mask extraction **(Fig. 1C)**. To evaluate the classifier, a model was trained on two of the three biological replicates and evaluated on its ability to classify individual worms between the two aforementioned groups on the remaining replicate for three total folds **(Fig. 1D)**.

This model achieved an AUC of 0.88 **(Fig. 1E),** confirming it was effective in distinguishing between the two classes and that a change in posture was strongly associated with exposure to a lethal heat shock with our assay conditions. **Fig. 1F** offers a visual demonstration of the model’s outputs, showing that the score is primarily quantifying how sinusoidal or straightened a worm is.

This model produces a probability that an individual worm has received a non-lethal or severe heat shock ranging from 0 to 1. By averaging these individual worm probabilities to the well level, we establish the Survival-Associated Posture score (SAP score), which we use for assay development, validation, and screening. Software used for this screen, broadly applicable to automatic analysis of *C. elegans* survival in microplate assays, is available as “ShapeScore” (https://github.com/tomh014/shape-score).

### Optimization of heat shock conditions

Consistent and effective heating of samples is a major source of variability in thermotolerance assays (*22*). To address this issue, we sandwiched 96-well plates between two one-centimeter-thick copper blocks pre-equilibrated to 35°C. We observed that plates heated by copper blocks in the incubator received a more effective heat shock as evidenced by lower survival at 10hrs of exposure compared with plates exposed to the air alone **(Fig. 2A)**. The “edge effect” is a common problem in microplate-based assays and can be caused by the tendency of edge wells to either heat up or evaporate faster than wells closer to the inside of the plate. As our assay uses impermeable plate seals, evaporation is likely not an issue. However, differences in heating between edge and interior wells still contribute to the edge effect. We found that survival was significantly lower in edge wells compared with central wells in plates heated only with air, and this was mitigated by the use of copper blocks **(Fig. 2B)**. This suggests that copper blocks not only deliver a more consistent heat shock, but also minimize edge effects, enabling us to confidently utilize the full plate for drug screening. All subsequent experiments therefore use the full 96-well plate with copper blocks.

**Figure 2.**
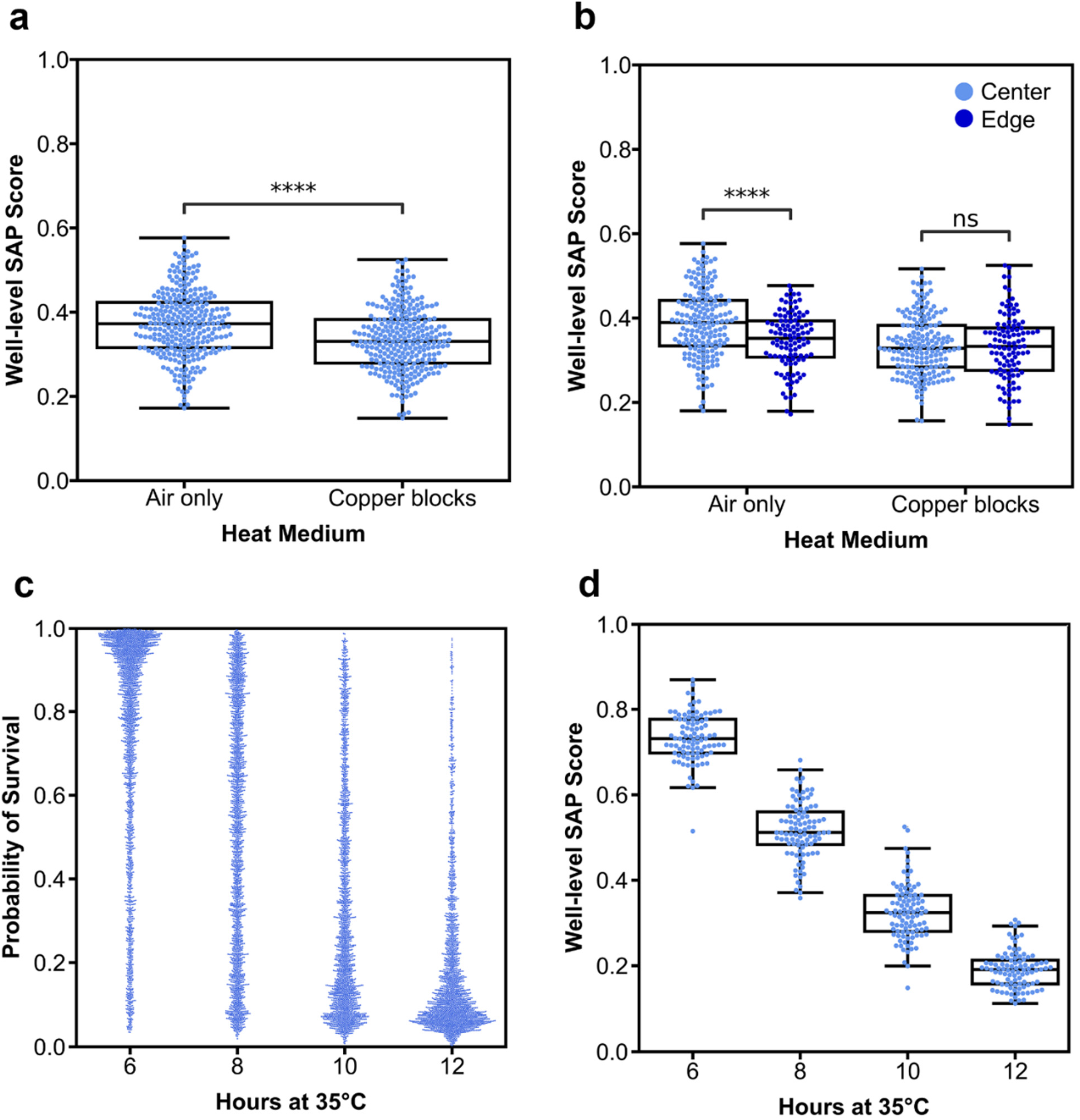
Survival-associated posture and differential heat exposure. (a) Comparison of well-level SAP score after 10hrs of heat exposure on plates placed in incubator versus plates sandwiched between copper blocks in incubator. (b) Comparison of edge (96-well plate rows A, H; cols 1, 12) and center wells in plates in incubator versus plates sandwiched between copper blocks in incubator. (c) Cross sectional time course with plates between copper blocks removed from incubator at specified timepoints (1 dot=1 worm mask). (d) Same experiment as (c) but summarized to the well level by averaging all individual mask probabilities in each well. Statistics: two-tailed t-tests, ns = not significant, **** p<0.0001.

We performed cross-sectional time course experiments to establish that the posture score is associated with the amount of time exposed to heat, and to find a timepoint that kills most control worms while still not being so lethal as to negatively influence the detection of hit compounds. **Fig. 2C** shows how individual worm survival probability distributions shift from mostly alive at 6hrs to mostly dead at 12hrs across four 96 well plates. Averaging individual worm probabilities within a well for a population-level summary survival score **(Fig. 2D)** shows a steady decrease in survival over time.

Bacterial concentration has also been shown to have effects on heat tolerance (*14*). We tested two concentrations (the higher one being the upper limit of what allowed clear images despite bacterial opacity): 2 and 4μL of 50x concentrated OP50. Accordingly, we observed increased heat tolerance with the higher concentration of OP50 **(Fig. S1)**. Since this indicates that worms are healthier when given more food, and the goal of the assay is to find interventions that allow healthy worms to mount a better stress response, we opted for 4μL.

To verify the ability of the assay to detect a known genetic intervention that increases heat resistance and lifespan, we compared wild-type N2 worms with worms harboring a mutation in the *age-1* gene that confers increased heat resistance (*2*). A time course experiment revealed separation of *age-1* and N2 SAP scores **(Fig. 3A, Fig. S2)** at multiple timepoints. To confirm that the assay can detect known chemical interventions that increase heat tolerance, we tested selected drugs (tetracycline, lipoic acid, and trehalose) that have been previously shown to increase stress resistance and lifespan (*6*, *23*) **(Fig. 3B, Fig. S2)**. Due to its efficacy, we used tetracycline to calibrate our assay and observed a strong separation from controls at 11hrs of heat exposure **(Fig. 3B)**. Calculation of the average precision score for each *age-1* and tetracycline biological replicate **(Fig. 3C)** verified that 11hrs provided the best separation between controls and both interventions out of the timepoints observed. We therefore performed our subsequent drug screen using 11hrs of heat exposure as our single-timepoint readout.

**Figure 3.**
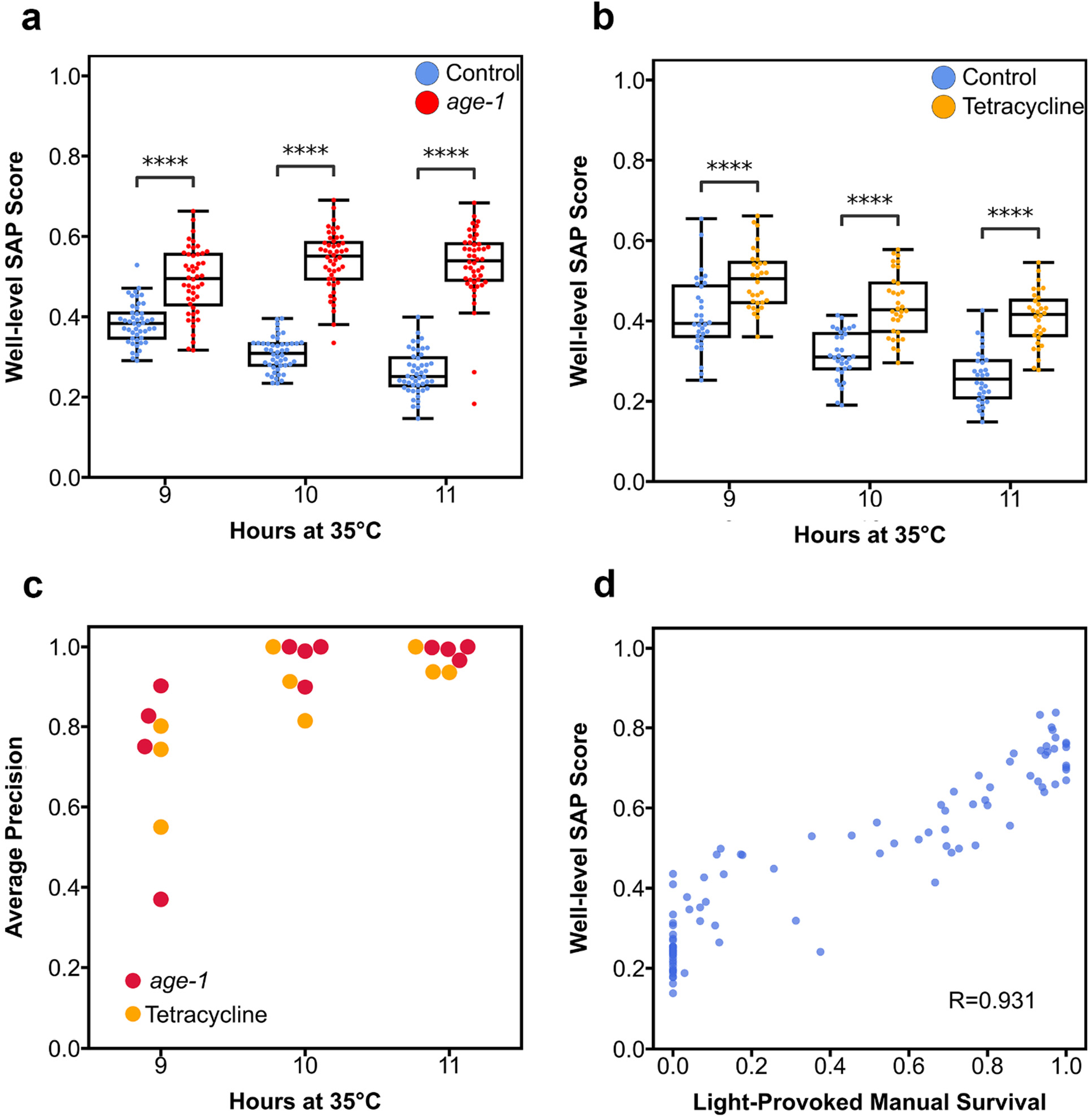
Validation of detection and survival measurement. (a) Well-level SAP score comparison of wild-type N2 worms versus age-1 mutant worms over time. (b) Well-level SAP score comparison of control (0.5% DMSO) worms versus tetracycline-treated (1mM) worms over time. (c) Average precision score of detection for each replicate at each timepoint, derived by treating controls wells as negative examples, colored by intervention. (d) Comparison of blue-light provoked movement-based survival versus the image-based automated SAP score (slope=0.468). Statistics: two-tailed t-tests, ns = not significant, **** p<0.0001.

We further validated the SAP score by comparing it to blue-light provoked movement, a previously established method of manually scoring worms for survival. At the various timepoints, specific wells were exposed to strong blue-light and then manually scored as alive or dead based on movement. The same plate was then exposed to UV (*24*) and imaged to generate automated posture scores that were compared with manual survival scores from their respective wells **(Fig. 3D)**. A significant Pearson correlation between the scores was observed (r=0.931, p=2.7×10^-27^). Of note is the variability of the SAP score in wells that are scored manually as having zero percent survival. This potentially reflects an ability of the posture score to assess worm health in more detail than a simple binary score but could also reflect insensitivity in detecting dead worms.

### A drug repurposing screen for heat stress resistance

The full screening workflow is illustrated in **Fig. 4A**. Worms are grown to adulthood on solid nematode growth media (NGM) plates with 24hrs of exposure to 50µM FUDR from the L4 stage. Day 1 adults are dispensed into 96 well plates in liquid media containing drug or DMSO (vehicle control) in the presence of FUDR. We opted to dispense 25 worms per well as a tradeoff between variability of dispensing and power of drug detection. After 24hrs of drug exposure, worms are subjected to an 11hr heat shock at 35°C which kills most DMSO-treated worms. After the 11hr heat shock, plates are returned to 20°C overnight, allowing worms that survived the heat shock time to recover, and critically injured worms time to die. Just prior to imaging, worms are subjected to brief UV stimulation to provoke movement in live worms and maximize the difference in posture compared with dead worms. The full well-plate is then imaged, and images are fed into the automated workflow.

**Figure 4.**
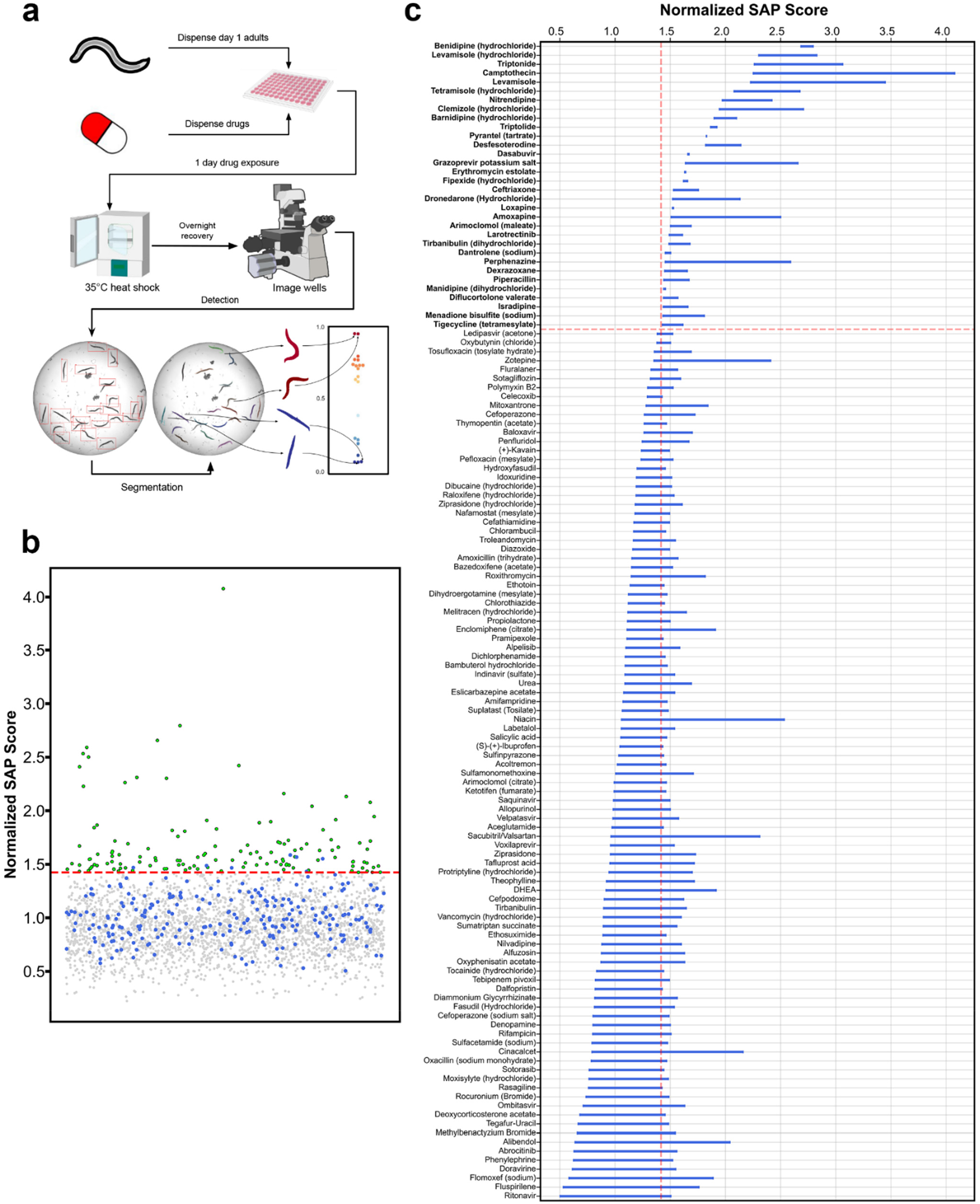
A screen for heat stress resistance with an FDA-approved drug library. (a) Drug screening workflow. (b) Control-normalized SAP scores for full library screen, with hits (green) called as drugs that confer scores two standard deviations above the mean (dotted line) of control wells (blue). All other non-hit compounds are gray. (c) Combined primary and secondary screen results for 130 drugs that passed primary screen. Each drug has two SAP scores from each screen, with the right side of the bar representing the higher score and the left representing the lower score. The vertical dotted line displays the two standard deviation cutoff and the horizontal line denotes chemicals that reached the cutoff.

We acquired a library of 2,782 FDA-approved drugs from MedChemExpress and screened a single replicate of each at 50µM for our primary screen **(Fig. 4B, Data S1)**. Eight 96-well plates were screened each week for a total of four weeks and this level of throughput was determined by the incubator size and the number of copper blocks available. Each plate contained eight control wells with DMSO vehicle alone. The SAP score of each drug was normalized to the means of control SAP scores for each plate. 130 primary hits were identified based on a cutoff of two standard deviations above the mean of all normalized control SAP scores. Rescreening the 130 primary hits at 50µM resulted in 31 final hits **(Fig. 4C)** using the same two standard deviation cutoff. We tested these final hits at two additional lower doses and found that all but one drug (triptolide) were most effective at the screening dose, 50µM **(Fig. S3)**.

### Identification of drugs that confer stress resistance and longevity

We next sought to determine which hits from the heat stress screen would also extend worm lifespan. We set up lifespans and transferred worms according to *Caenorhabditis* Interventions Testing Program protocols (*25*). To expedite the screening of the 31 hits from the screen, we used a single timepoint assay in which worms were transferred to new drug plates every two to three days but were only scored for survival on day 12. Plates were prepared with 0.5% DMSO and 50µM FUDR and drug treatment began on day one of adulthood. This shortened assay enabled us to screen all 31 candidate drugs at two doses (5µM, 50µM) in triplicate in six weeks.

Overall, we found that 18/31 candidates nominated from the heat stress assay extended lifespan at one or two doses, with 16/31 conferring a significant increase in day 12 survival at 50µM **(Fig. 5A, Data S2)**. Two compounds only extended lifespan at the 5µM dose (dexrazoxane and diflucortolone). Only four of the compounds that extend lifespan have previously been reported to be geroprotective (nitrendipine, amoxapine, loxapine (*7*), and triptolide (*26*)). We selected the six compounds with the strongest effects on day 12 for analysis in longitudinal lifespans: tigecycline, ceftriaxone, clemizole, pipericillin, isradipine and camptothecin **(Fig. 5B-G)**. All six drugs conferred significant increases in lifespan (Cox proportional hazard, p<0.05) at 50µM for at least 2/3 biological replicates **(Fig. S4, Data S3)**, demonstrating both the efficacy of the single timepoint lifespan assay and the heat shock assay for nominating candidates for life extension. Three drugs, ceftriaxone, tigecycline, and isradipine, conferred mean increases in lifespan over 30% **(Table S1)**.

**Figure 5.**
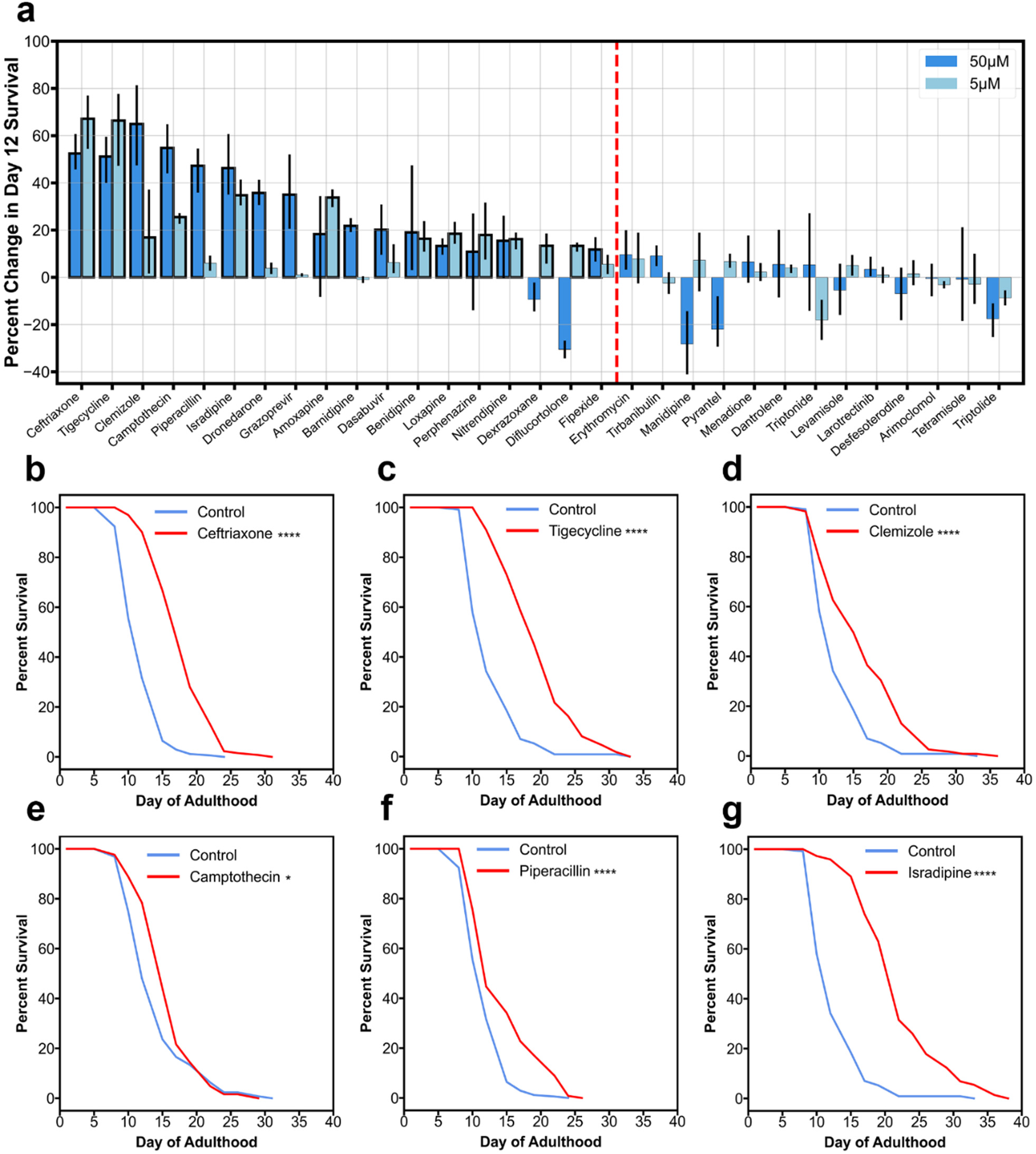
Chemicals that increase heat stress resistance increase survival. (a) Single-timepoint (D12 of adulthood) survival quantification of 31 hit compounds on standard NGM compared to controls, performed in triplicate (SE bars displayed). Bold bars indicate statistical significance (one-sided, two-sample independent proportion test, FDR<0.05). Drugs to the left of the dotted red line were significant at one or two of the tested doses. (b-g) Longitudinal lifespans of ceftriaxone, tigecycline tetramesylate, clemizole, camptothecin, piperacillin, and isradipine at 50μM. Cox-proportional hazards test for significance (* p<0.05, **** p<0.00005).

### *In vitro* senotherapeutic activity of life-extending drugs

To investigate whether our drugs that extend *C. elegans* lifespan could translate to a human cell-based model of aging, we assayed the top six lifespan hits for their ability to reduce cellular senescence in human umbilical vein endothelial cells (HUVECs) induced to senescence by oxidative stress (H_2_O_2_). Senescent and non-senescent (proliferating) cells were treated with tigecycline, ceftriaxone, clemizole, piperacillin, isradipine or camptothecin over a 24hr period, after which a fluorogenic dye, C_12_FDG, was used to stain SA-β-gal-positive cells (*27*). Clemizole, isradipine and piperacillin displayed a significant senomorphic effect in senescent HUVECs, effectively reducing the relative number of SA-β-gal-positive cells without inducing total cell loss and without detectable effects in proliferating cells **(Fig. 6)**. No senotherapeutic effects were observed after tigecycline, ceftriaxone and camptothecin treatment **(Fig. S5).**

**Figure 6.**
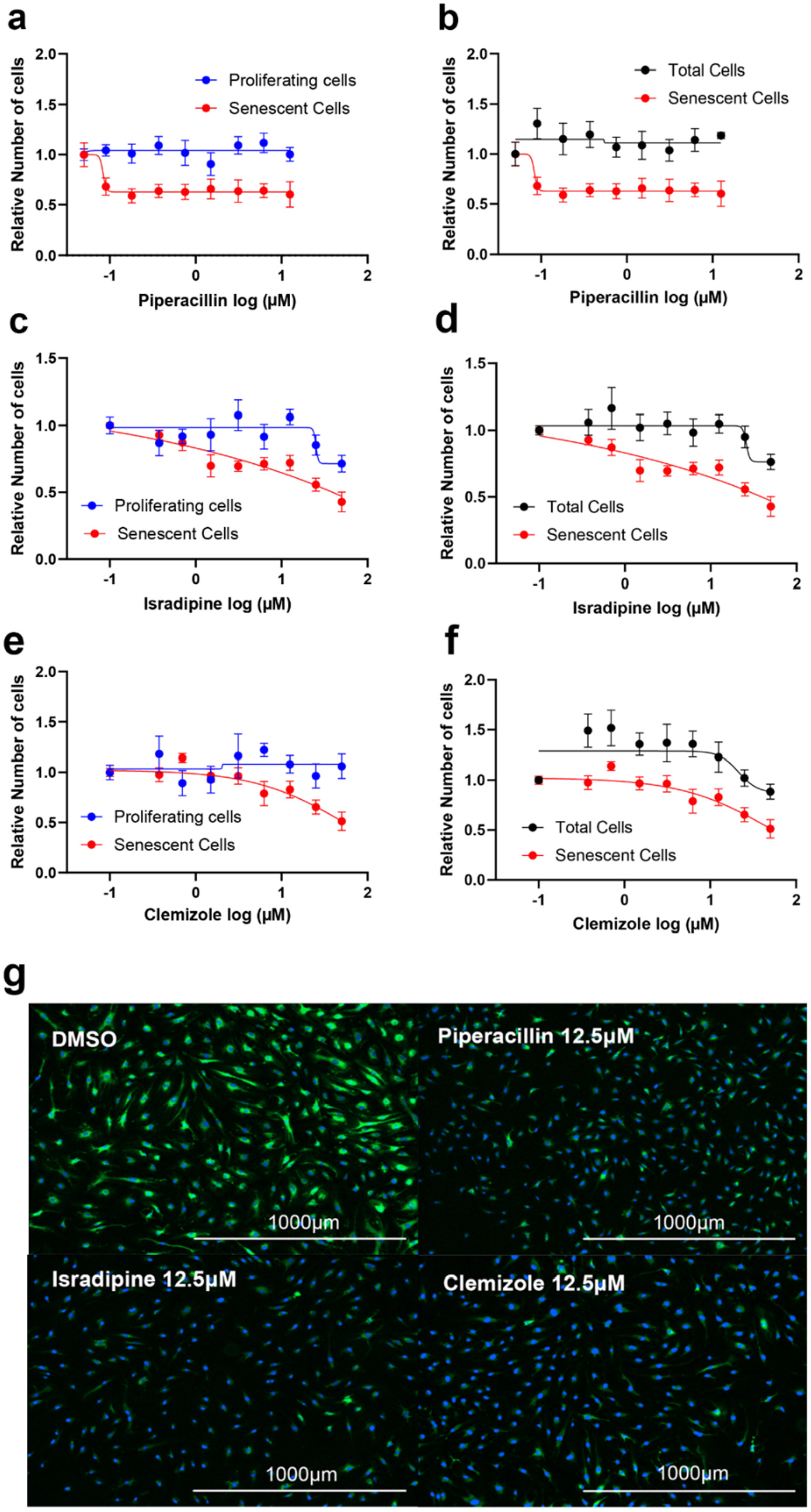
Selected life-extending chemicals modulate senescence *in vitro*. (a-c) Comparison of proliferating, nonsenescent cells and senescent cells (HUVECs) after induction of senescence by exposure to hydrogen peroxide with different drug treatments. (d-f) Comparison of total cells to senescent cells with different drug treatments. (g-j) Nuclear (blue) and senescent (green) stained cells treated with different drugs after induction of senescence. Blue stained cells that are not stained green represent nonsenescent cells. Error bars denote standard error of the mean.

## Discussion

Since the discovery that single gene mutations in *C. elegans* extend lifespan through evolutionarily conserved mechanisms, there has been interest in using the nematode to identify drugs that target aging and improve healthspan (*28*). Several approaches for improving throughput have been developed, but scalability and ease of implementation remain a challenge for *C. elegans* phenotypic screens for survival. We present a method that uses off-the-shelf equipment for dispensing worms into 96-well plates and an analysis pipeline that is based on low-magnification bright-field images. By using heat stress resistance as the primary screening phenotype, the assay can be completed in less than three days, compared with two to five weeks for screens that focus on lifespan. In this study, the limiting factors to throughput were the incubator size and the number of copper blocks, two issues that could easily be overcome. Therefore, with a relatively small investment, this assay could be comfortably scaled to screen several thousand compounds per week.

Despite the short length of our assay, we included FUDR in our protocol to limit progeny production. This not only helps maintain the appropriate food concentration over the course of the 2.5 days that worms are in liquid culture but also helps maintain image quality for the analysis step. FUDR is commonly used in *C. elegans* aging studies but some reports have suggested that it can interact with certain genetic backgrounds to affect lifespan (*29*). It is therefore feasible that there could also be some compound-specific interactions with FUDR that impact survival. However, since follow-up lifespans are likely to be performed in the presence of FUDR for ease of throughput, we felt that the potential problems were outweighed by the benefits. Future assays could employ alternate methods of progeny control such as an auxin-induced genetic system for induction of sterility (*30*).

We performed our primary screen with one biological replicate and only 31 out of the 130 primary hits passed the cutoff upon retesting, consistent with known variability issues in thermotolerance assays (*22*). By moving forward only with drugs that passed the cutoff twice, we enriched our hits for drugs that strongly increase heat stress resistance, with the caveat that we have an unknown but likely appreciable false negative rate, which is a common feature of high-throughput screens. Drugs that conferred high increases in heat tolerance for the primary screen but narrowly missed the cutoff for the rescreen could warrant further investigation. Zotepine, for example **(Fig. 4C, rank 35)**, has previously been shown to strongly increase lifespan in liquid media (*31*), and belongs to the same class as two compounds that passed the rescreen and extended lifespan in the single timepoint assay, amoxapine and loxapine (ranks 18 and 19). If chemical quantity is not a limiting factor, we would advocate duplicate or triplicate primary screens. However, our data show that, with a single primary screen and focused retesting, we can identify compounds that increase lifespan.

Our library had an overlap of about 348 chemicals with the LOPAC 1280 library used in a previous screen for lifespan (*7*), 24 of which were found to extend lifespan using only a single drug dose. Three of these drugs (amoxapine, loxapine, and nitrendipine) were hits in our heat stress assay and extended lifespan in the single time point assay. Conversely, three of our lifespan hits (perphenazine, camptothecin, and clemizole) were not detected in the liquid lifespan assay. Another hit from our assay, tigecycline did not extend lifespan in liquid (*32*), but extended lifespan robustly in our solid-media assay **(Fig. 5C)**. Reasons for these differences may include dose, timing and frequency of drug treatments. *C. elegans* has a robust drug metabolism system (*33*) and thus it is possible that single drug exposures at the beginning of lifespan experiments are associated with loss of bioactivity as the experiment progresses. The short length of the heat shock assay potentially mitigates this problem as worms are subjected to heat stress within 24hrs of exposure to the compounds. The differences in hits might also generally reflect differences in heat stress versus lifespan as longevity-associated phenotypes. In addition, the effect of compounds on nematode lifespan is well-known to be highly dependent on assay and lab-specific conditions. This has been demonstrated by the *Caenorhabditis* Interventions Testing Program (CITP), which is focused on characterizing drugs that extend lifespan in worms with diverse genetic backgrounds (*34*).

We identified multiple drugs that have not previously been associated with longevity. Ceftriaxone is a cephalosporin antibiotic that has demonstrated neuroprotective effects in a mouse neurodegenerative disease model (*35*). Tigecycline is tetracycline antibiotic and multiple members of this class have previously been shown to extend lifespan (*7*, *36*), potentially acting through attenuation of translation (*32*).

Clemizole is serotonin receptor agonist and a putative therapeutic for Dravet Syndrome, a form of childhood epilepsy (*37*). Serotonin-modulating antidepressants have previously been demonstrated to positively affect longevity (*5*). Several hits are members of shared classes which have both been previously characterized as geroprotective (*5*, *7*). These included tricyclic antidepressants amoxapine and loxapine, and multiple dihydropyridine calcium channel blockers (isradipine, nitrendipine, benidipine, and barnidipine).

A long-term goal of drug screens in worms is to identify compounds that have translational utility in slowing aging in humans. Compounds that slow aging in multiple model species are more likely to act through conserved mechanisms, but mouse longevity studies are lengthy and expensive. An alternative approach is to examine their effects on human cellular senescence. In general, the overlap between senotherapeutic drugs and drugs that extend life in non-mammalian model animals is not well-studied, but there are multiple examples of drugs that extend lifespan in *C. elegans* and act as senomodulators *in vitro* including metformin and rapamycin (*38*), two of the most prominent putative geroprotective drugs targeting aging in the field.

From our screen, we found that three of the chemicals that extend lifespan in *C. elegans* also possess potential senomorphic activity in human endothelial cells. Senomorphics are senotherapeutic drugs that do not necessarily kill senescent cells, but prevent their progression and development of the SASP (senescence-associated secretory phenotype), as opposed to senolytics, which selectively kill senescent cells (*38*). We identified isradipine, a calcium channel blocker, as potentially senomorphic. A related compound, barnidipine, was previously found to be senolytic in a model of cigarette-smoke induced senescence (*39*). The prevalence of calcium channel blockers in our screen hits, as well as another screen for life extension in worms (*7*) suggests that calcium signaling is an important, conserved molecular mechanism linking senescence and longevity. Although the data are intriguing, these drugs could be extending life through a number of different mechanisms, and further investigation into the connection between senotherapeutics and nematode life extenders is warranted.

The utility of *C. elegans* as a system for high-throughput screening is not limited to geroscience (*40*). Our machine learning approach could be readily adapted to improve throughput and scoring in other survival based assays such as hypoxic stress (*41*), as well as screens for anthelmintics (*42*, *43*). In this latter respect, we identified multiple drugs that reduce worm survival including a known anthelmintic, niclosamide (*44*) **(Data S1).** Many other compounds that reduce survival are not currently used as anthelmintics, indicating potential new candidates. Moreover, with appropriate training data, our system could be adapted to other posture-based phenotypes such as paralysis, enabling high-throughput screens in worm models of neurodegenerative disease (*45*).

As the field of geroscience matures, there will be an increasing need to identify compounds that have therapeutic potential for increasing human healthspan (*28*). The nematode *C. elegans* has been instrumental in defining genetic and chemical interventions that impact longevity. With this project, we have developed an automated and scalable assay to nominate drug candidates for geroprotection, leading to new compounds that not only extend lifespan in worms but also translate to a human model of aging. The discovery of multiple drugs that are effective in these established models of longevity underlines the importance of developing new phenotypic assays to establish a more diverse pool of candidate drugs. In addition, the automated nature of the readout facilitates scaling of these assays, which could lead to more effective candidate discovery in the future.

## Materials and Methods

### Chemicals

A library of 2,782 FDA-approved drugs was obtained from MedChemExpress. Drugs were distributed into flat-bottom 96-well tissue culture plates from V-bottom 96-well source plates.

The following drugs/salts were ordered for lifespan experiments: grazoprevir potassium salt, tirbanibulin dihydrochloride, desfesoterodine (MedChemExpress); dronedarone hydrochloride (Tci America); larotrectinib (Apexbio); tetramisole hydrochloride (Merck Millipore); erythromycin estolate, ceftriaxone sodium, barnidipine hydrochloride, fipexide hydrochloride, pyrantel tartarate, dantrolene sodium salt, tigecycline tetramesylate, diflucortolone valerate, loxapine (AstaTech); benidipine hydrochloride, piperacillin sodium salt, triptolide, menadione sodium bisulfite, isradipine, (S)-(+)-camptothecin, dexrazoxane (Chem Impex); perphenazine (Oakwood Products), clemizole hydrochloride (Cayman); amoxapine (Sigma-Aldrich); manidipine dihydrochloride (G Biosciences); dasabuvir, arimoclomol maleate, nitrendipine (Ambeed); triptonide (TARGETMOL).

### Strains and maintenance

*C. elegans* strains were maintained as previously described (*46*). Bristol N2 (wild-type) and TJ1052[*age-1(hx546)*] were obtained from the *Caenorhabditis* Genetics Center (University of Minnesota, MN). Worms were maintained at 20°C unless otherwise stated. For mass cultures and the heat shock assay, concentrated *E. coli* OP50 was used. An overnight culture was concentrated by centrifugation at 5000g for 15 minutes in 500mL centrifuge tubes and pellets were resuspended in S-basal at 1/50th the original volume. 4µL of 50x OP50 corresponds to OD600 ∼ 1 when diluted in 100µL S-basal.

### Heat shock assay

Worm mass cultures used 10cm petri dishes with 25mL of NGM. 40 eggs were placed on one 10cm plate seeded with 2mL of 50x concentrated OP50. One week later, eggs were obtained by bleach treatment (*47*) and hatched into S-basal to generate synchronized L1s. 24hrs later, 20,000 L1 worms were seeded onto a 10cm plate with 2mL of 50x OP50 and grown at 20°C for until the L4 stage (44hrs). L4s were then transferred to a 10cm plate containing 3mL of 50x OP50 and 50µM FUDR and grown for an additional 24hrs. D1 adults were recovered from the FUDR plate, washed with one volume of S-basal wash and resuspended in S-basal containing 50µM FUDR, 4% 50x OP50 to a concentration of 25 worms/100µL in a glass container with a magnetic stir bar. Worms were dispensed (100µL) using an Agilent MultiFlo liquid dispenser with a wide bore dispense cassette into plates containing DMSO vehicle or drug such that the final treatment concentration was 0.5% DMSO unless otherwise noted. Plates were then sealed with standard impermeable PCR plate seals, and worms were incubated with drug for 24hrs at 20°C before the heat shock. The next day, worms were sandwiched between copper blocks that had been equilibrating at 35°C in an EchoTherm (Torrey Pines Scientific) incubator for a minimum of three days. Plates were removed at defined timepoints and allowed to rest overnight at 20°C (all 26hrs after the beginning of the heat shock). The following steps happened plate by plate, the next day. A 96 well plate was shaken on an orbital plate shaker at max settings for 20s after removing the seal. The MultiFlo was used to dispense 230µL of S-basal into the entire plate such that the total volume was flush with the top of the well and the meniscus was minimized. The plate was then placed on a standard UV transilluminator for 2min and then allowed to rest for 2min. Plates were then imaged at 2x magnification with the brightfield channel using a stage-motorized Nikon Ti2 Eclipse. Plates were discarded after imaging.

### Single-timepoint lifespan assays

Synchronous egg populations were obtained by allowing 20 day 1 adults to lay eggs over a 4hr period. 100 synchronized day 1 adults from the egg lay were moved onto triplicate drug plates (50µM or 5µM drug, 0.5% DMSO, 50µM FUDR). FUDR and drug/DMSO were dripped onto the 10mL plate to achieve the desired concentrations. Worms were transferred on D2, D5, D8, D10, and manually scored on D12. All scores were made blind to the treatment.

### Longitudinal lifespan assays

The setup for longitudinal assays was identical to single timepoint assays, but with two technical replicates of 75 worms for each condition, for each bioreplicate. Worms were transferred on days 2, 5, 8, 10, 12, 15, 19, 22, and weekly after day 22 until all worms were dead. Survival was scored every 2-3 days by touch-provoked movement.

### Image analysis

Images were saved as ome.tiff files and converted to jpeg format (unbounded tiff intensity was normalized by the maximum intensity detected in each experiment for jpeg conversion). Faster R-CNN (using a finetuned Resnet18 backbone) was used for bounding box generation on jpeg well images. Bounding boxes were used as prompts for SAM 2 (sam2_hiera_tiny.pt). The top mask prediction from each bounding box region was extracted for each well and used for downstream analysis after a size filter (only masks with a minimum length/width above 50 pixels and a maximum length/width below 600 pixels are kept). Masks were made a single color by setting nonzero integer values to 255 using Pillow and Numpy. The binary dead/alive classifier was run on the unicolor masks to derive individual worm survival probabilities. General analysis of data and figure generation was achieved using various Python packages including Numpy, Matplotlib, Seaborn, statsannotations, SciPy, statsmodels, and Pandas. More details can be found in freely available code at https://doi.org/10.5281/zenodo.20437182. Disclosure: generative AI tools (Gemini 3.1 Pro and ChatGPT GPT-4o) were used to assist in the creation of this software.

### Model training

All survival classifiers were trained on binarized segmentation pipeline outputs. The final deployment classifier was trained on 16,226 worms treated with a short, non-lethal heat shock (4hrs, 6hrs) and 16,335 worms treated with a long, severe heat shock (10hrs, 12hrs, 14hrs). Models were finetuned Resnet18 backbones downloaded from Pytorch’s Model Zoo. Models were trained for 25 epochs using an SGD optimizer with learning rate=0.001, momentum=0.9, cross entropy loss, and stepLR scheduler with a step size=7 and gamma=0.1 (*48*). All datasets and classifier models can be generated from available code.

### Evaluation of classifiers and detection

To evaluate the survival classifier, individual worm masks were given binary labels based on the amount of time exposed to heat (based on exposure to a nonlethal or severe heat shock). Classifiers were trained on two biological replicates (6hrs, 10hrs, 12hrs) supplemented with additional experiments (4hrs, 14hrs) and evaluated on the remaining replicate (6hrs, 10hrs, 12hrs) for three folds. Test fold predictions and labels were concatenated to produce the ROC curve using sklearn. Exact projects and split construction details can be found in code (CROSSVAL1_make_dataset_binary.py). 4 and 6hrs were defined as live, and 10, 12, and 14hrs were defined as dead. To evaluate detection of interventions **(Fig. 3C)**, well-level SAP scores were assigned ground truth labels based on perturbation (control as a negative example, tetracycline or *age-1* as positive examples). Average precision score was calculated using sklearn.

### Senotherapeutic drug screening

Human Umbilical Vein Endothelial Cells (HUVECs) were cultured in Endothelial cell growth base media (R&D, CCM027 and CCM029) containing 1% Penicillin / Streptomycin. HUVECs were induced to undergo senescence by treatment with 100 µM hydrogen peroxide (H2O2) for 3hrs during 3 consecutive days, followed by 3 days in normal culture media. SA-β-gal activity was tested using a C_12_FDG staining assay. HUVECs were plated in 96-well black wall clear bottom plates (Corning, 3603) in triplicate at a density of 2500 cells/well. Following the addition of drugs (isradipine, clemizole hydrochloride, piperacillin sodium, ceftriaxone sodium, tigecycline tetramesylate, (S)-(+)-Camptothecin) or vehicle, the cells were incubated for 24hrs at 37℃ and 5% CO2. After removing the medium, cells were incubated with 100nM bafilomycin A1 (Selleckchem, S1413) in normal culture medium for 1hr to induce lysosomal alkalinization, followed by incubation with 20µM fluorogenic substrate C12FDG (Cayman Chemical, 25583) for 2hrs and counterstaining with 2µg/ml Hoechst 33342 (Invitrogen, H1399; 1:2000 dilution) for 15 minutes. Finally, the cells were washed with PBS and immediately imaged in six fields per well using the high-content fluorescence image acquisition and analysis platform Cytation 1 (BioTek).

## Acknowledgments

Some strains were provided by the *C. elegans* Genetics Center (CGC), which is funded by NIH Office of Research Infrastructure Programs (P40 OD010440). Thanks to Dr. Gordon Lithgow and David Hall for constructive feedback over the course of the development of the screen. Thanks to the Minnesota Supercomputing Institute for high-performance computing resources and storage during the development of the project, and to Dr. Ham C. Lam for high-performance computing resource guidance.

## Data availability

All raw images used to produce figures for all validation and screening experiments are provided in heatshock.zip at https://doi.org/10.5281/zenodo.20437182, along with code/models to produce masks and predictions. Pre-ran worm mask predictions for each project are available in each project folder. Code to reproduce all figures from raw data is available in figures.zip. Screen results (including normalized scores for heat stress resistance, declared hits, and chemical structures) and lifespan data are available as Extended Data. A user-friendly Python library, along with tutorials, is available as ShapeScore https://github.com/tomh014/shape-score.

## Supplementary Figures

**Fig. S1.**
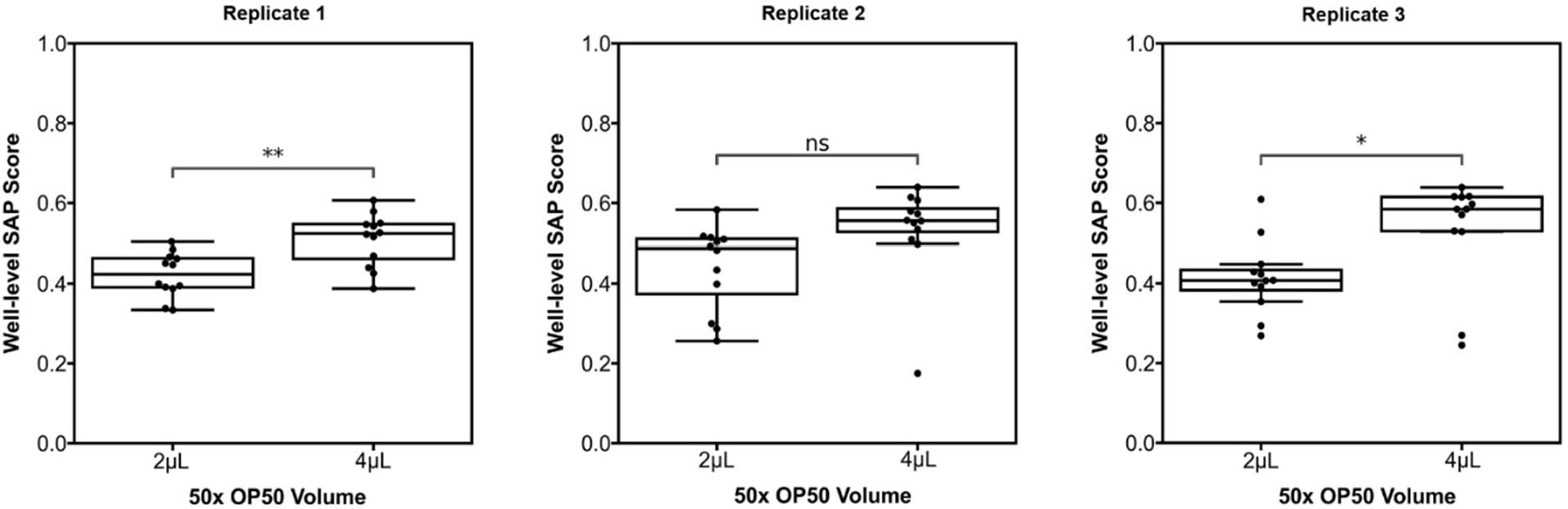
Comparison of SAP scores after a 9hr heat shock at 35°C from worms fed 4 or 2 μL of 50x concentrated OP50. (** p<0.01, * p<0.05, ns p>=0.05, t-test)

**Fig. S2.**
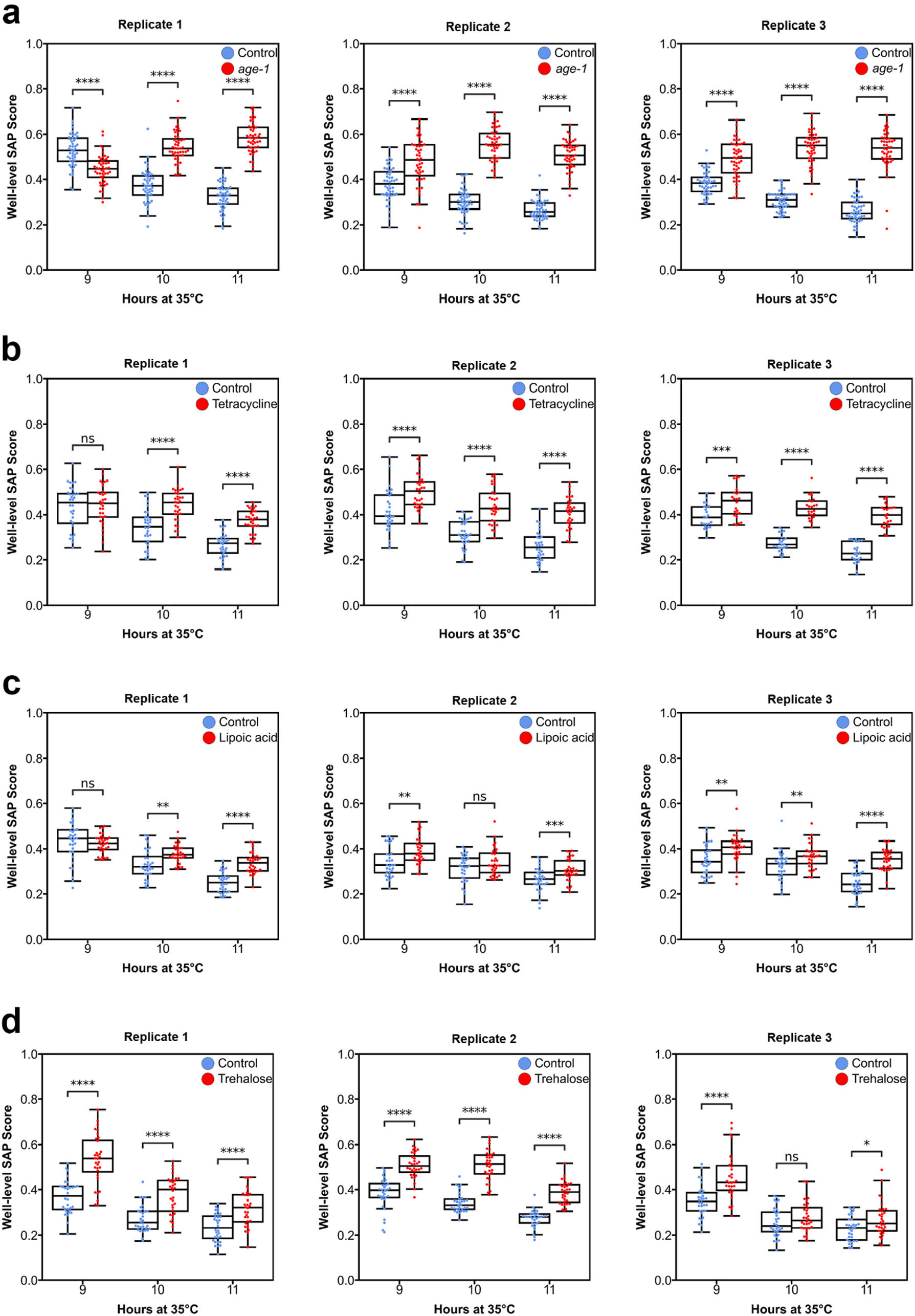
Timecourse experiments showing SAP scores of varied interventions compared to controls after heat shock from (a) age-1 worms, (b) 1mM tetracycline, (c) 1mM lipoic acid, and (d) 1mM trehalose. (**** p<0.0001, ****p<0.001, ** p<0.01, * p<0.05, ns p>=0.05, t-test).

**Fig. S3.**
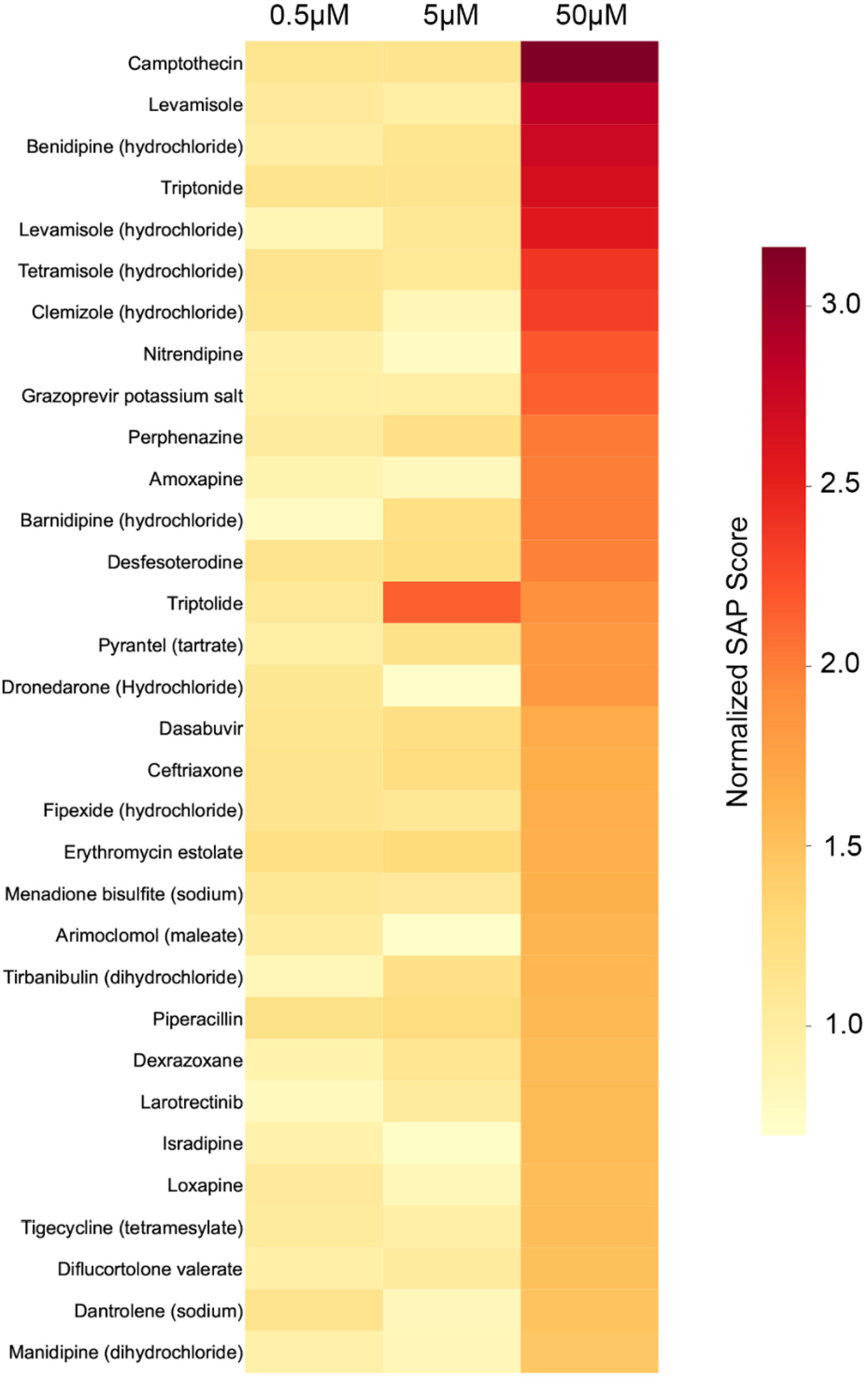
Dose response of hit drugs. 31 hit compounds retested at varying doses in the heat shock assay. SAP scores represent two-replicate averages.

**Fig. S4.**
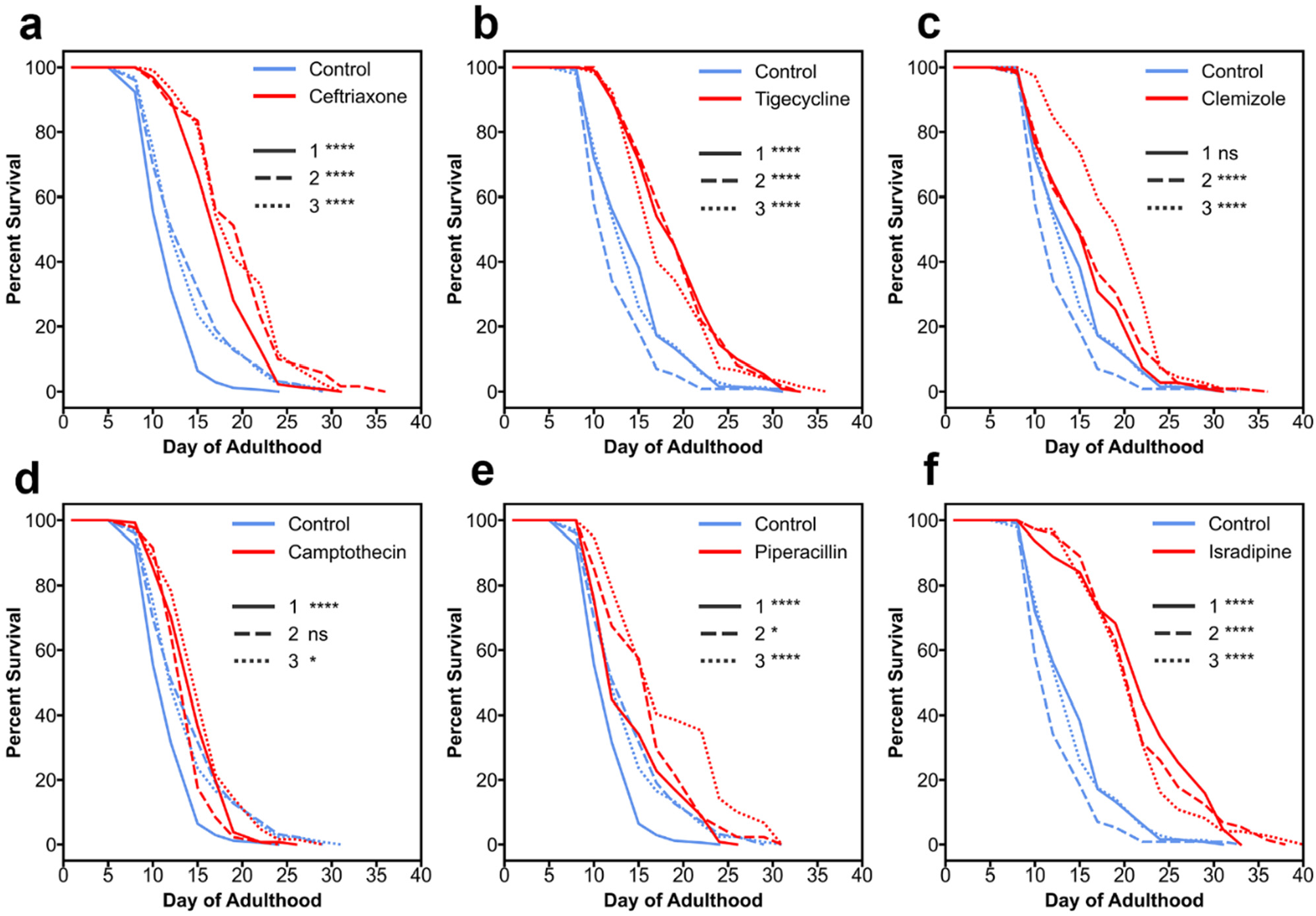
Longitudinal lifespan experiments. (a-f) Survival curves for all drugs. Each dotted line represents a separate biological replicate. Cox-proportional hazards test for significance (ns = not significant, * p<0.05, **** p<0.00005).

**Fig. S5.**
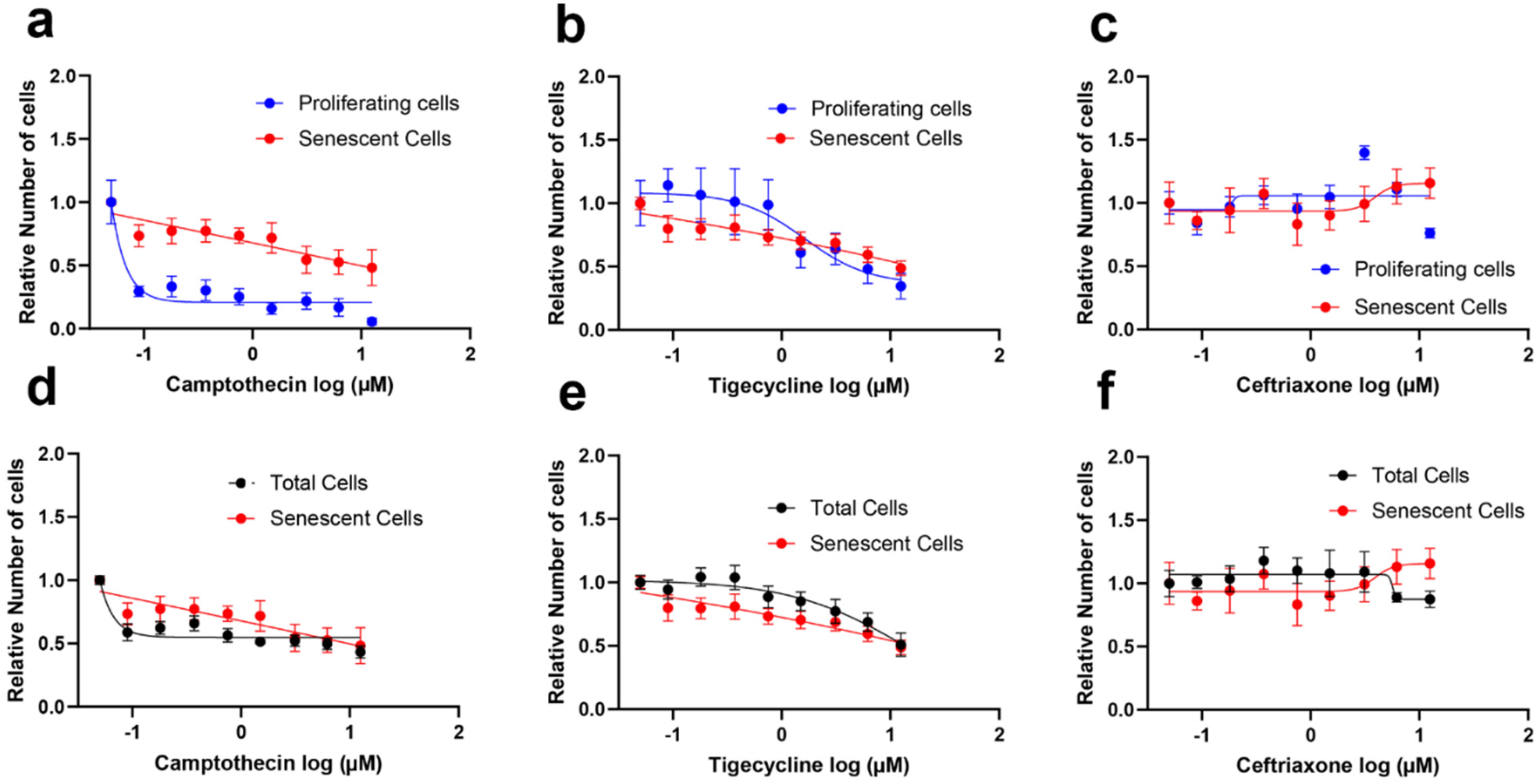
Life-extending drugs that are not protective in senescent HUVECs. (a-c) senescent cells versus proliferating cells. (d-f) senescent cells versus total cells. Error bars denote standard error of the mean.

**Table S1.** Heat stress hits that extend longitudinal lifespan. Column: 1) Drug name. 2) Target category provided by MedChemExpress. 3) Class provided by MedChemExpress. 4) Percent increase in mean lifespan of all recorded worms for an intervention. 5) Drug formulation molecular weight.

| Name | Target | Class | % increase in mean lifespan |
| --- | --- | --- | --- |
| Ceftriaxone sodium | Antibiotic; Aurora Kinase; Bacterial; GSK-3 | Anti-infection; Cell Cycle/DNA Damage; Epigenetics; PI3K/Akt/mTOR; Stem Cell/Wnt | 43 |
| Tigecycline tetramesylate | Antibiotic; Autophagy; Bacterial | Anti-infection; Autophagy | 36.9 |
| Clemizole hydrochloride | HCV; HCV Protease; Histamine Receptor; TRP Channel | Anti-infection; GPCR/G Protein; Immunology/Inflammation; Membrane Transporter/Ion Channel; Metabolic Enzyme/Protease; Neuronal Signaling | 23 |
| (S)-(+)-Camptothecin | ADC Payload; Antibiotic; Apoptosis; Fungal; Influenza Virus; MicroRNA; Topoisomerase | Antibody-drug Conjugate/ADC Related; Anti-infection; Apoptosis; Cell Cycle/DNA Damage; Epigenetics | 12 |
| Piperacillin sodium salt | Antibiotic; Bacterial; Beta-lactamase; Penicillin-binding protein (PBP) | Anti-infection | 23.2 |
| Isradipine | Autophagy; Calcium Channel | Autophagy; Membrane Transporter/Ion Channel; Neuronal Signaling | 55.3 |

## Extended Data

**Data S1.** Heat stress screen results for 2,782 FDA-approved drug library.

**Data S2.** Single-timepoint lifespan assay survival data for 31 final hit compounds.

**Data S3.** Longitudinal lifespan data for six top hits.

## References

1. M. R. Klass, A method for the isolation of longevity mutants in the nematode Caenorhabditis elegans and initial results. Mechanisms of Ageing and Development 22, 279–286 (1983).

2. D. B. Friedman, T. E. Johnson, A mutation in the age-1 gene in Caenorhabditis elegans lengthens life and reduces hermaphrodite fertility. Genetics 118, 75–86 (1988).

3. B. Hamilton, Y. Dong, M. Shindo, W. Liu, I. Odell, G. Ruvkun, S. S. Lee, A systematic RNAi screen for longevity genes in *C. elegans*. Genes Dev. 19, 1544–1555 (2005).

4. M. Hansen, A.-L. Hsu, A. Dillin, C. Kenyon, New Genes Tied to Endocrine, Metabolic, and Dietary Regulation of Lifespan from a Caenorhabditis elegans Genomic RNAi Screen. PLoS Genet 1, e17 (2005).

5. M. Petrascheck, X. Ye, L. B. Buck, An antidepressant that extends lifespan in adult Caenorhabditis elegans. Nature 450, 553–556 (2007).

6. M. G. Benedetti, A. L. Foster, M. C. Vantipalli, M. P. White, J. N. Sampayo, M. S. Gill, A. Olsen, G. J. Lithgow, Compounds that confer thermal stress resistance and extended lifespan. Experimental Gerontology 43, 882–891 (2008).

7. X. Ye, J. M. Linton, N. J. Schork, L. B. Buck, M. Petrascheck, A pharmacological network for lifespan extension in *C aenorhabditis elegans*. Aging Cell 13, 206–215 (2014).

8. M. Lucanic, T. Garrett, I. Yu, F. Calahorro, A. Asadi Shahmirzadi, A. Miller, M. S. Gill, R. E. Hughes, L. Holden-Dye, G. J. Lithgow, Chemical activation of a food deprivation signal extends lifespan. Aging Cell 15, 832–841 (2016).

9. B. K. Kennedy, S. L. Berger, A. Brunet, J. Campisi, A. M. Cuervo, E. S. Epel, C. Franceschi, G. J. Lithgow, R. I. Morimoto, J. E. Pessin, T. A. Rando, A. Richardson, E. E. Schadt, T. Wyss-Coray, F. Sierra, Geroscience: Linking Aging to Chronic Disease. Cell 159, 709–713 (2014).

10. G. J. Lithgow, T. M. White, S. Melov, T. E. Johnson, Thermotolerance and extended life-span conferred by single-gene mutations and induced by thermal stress. Proc. Natl. Acad. Sci. U.S.A. 92, 7540–7544 (1995).

11. M. J. Muñoz, D. L. Riddle, Positive Selection of *Caenorhabditis elegans* Mutants With Increased Stress Resistance and Longevity. Genetics 163, 171–180 (2003).

12. H. Ji, D. Chen, C. Fang-Yen, Automated multimodal imaging of *Caenorhabditis elegans* behavior in multi-well plates. GENETICS, iyae158 (2024).

13. N. Stroustrup, B. E. Ulmschneider, Z. M. Nash, I. F. López-Moyado, J. Apfeld, W. Fontana, The Caenorhabditis elegans Lifespan Machine. Nat Methods 10, 665–670 (2013).

14. M. S. Gill, A. Olsen, J. N. Sampayo, G. J. Lithgow, An automated high-throughput assay for survival of the nematode Caenorhabditis elegans. Free Radical Biology and Medicine 35, 558–565 (2003).

15. T. I. Moy, A. L. Conery, J. Larkins-Ford, G. Wu, R. Mazitschek, G. Casadei, K. Lewis, A. E. Carpenter, F. M. Ausubel, High-Throughput Screen for Novel Antimicrobials using a Whole Animal Infection Model. ACS Chem. Biol. 4, 527–533 (2009).

16. K. He, X. Zhang, S. Ren, J. Sun, Deep Residual Learning for Image Recognition. arXiv arXiv:1512.03385 [Preprint] (2015). 10.48550/arXiv.1512.03385.

17. A. Krizhevsky, I. Sutskever, G. E. Hinton, ImageNet classification with deep convolutional neural networks. Commun. ACM 60, 84–90 (2017).

18. M. Deserno, K. Bozek, WormSwin: Instance segmentation of C. elegans using vision transformer. Sci Rep 13, 11021 (2023).

19. P. E. L. Castro, K. Kounakis, A. G. Garví, I. Gkikas, I. Tsiamantas, N. Tavernarakis, A.-J. Sánchez-Salmerón, SegElegans: Instance segmentation using dual convolutional recurrent neural network decoder in Caenorhabditis elegans microscopic images. Computers in Biology and Medicine 190, 110012 (2025).

20. S. Ren, K. He, R. Girshick, J. Sun, Faster R-CNN: Towards Real-Time Object Detection with Region Proposal Networks. IEEE Trans. Pattern Anal. Mach. Intell. 39, 1137–1149 (2017).

21. N. Ravi, V. Gabeur, Y.-T. Hu, R. Hu, C. Ryali, T. Ma, H. Khedr, R. Rädle, C. Rolland, L. Gustafson, E. Mintun, J. Pan, K. V. Alwala, N. Carion, C.-Y. Wu, R. Girshick, P. Dollár, C. Feichtenhofer, SAM 2: Segment Anything in Images and Videos. arXiv arXiv:2408.00714 [Preprint] (2024). 10.48550/arXiv.2408.00714.

22. S. C. Zevian, J. L. Yanowitz, Methodological considerations for heat shock of the nematode Caenorhabditis elegans. Methods 68, 450–457 (2014).

23. D. B. Lombard, W. J. Kohler, A. H. Guo, C. Gendron, M. Han, W. Ding, Y. Lyu, T.-T. Ching, F.-Y. Wang, T. S. Chakraborty, Z. Nikolovska-Coleska, Y. Duan, T. Girke, A.-L. Hsu, S. D. Pletcher, R. A. Miller, High-throughput small molecule screening reveals Nrf2-dependent and -independent pathways of cellular stress resistance. SCIENCE ADVANCES (2020).

24. S. K. Rajasekharan, C. J. Raorane, J. Lee, LED based real-time survival bioassays for nematode research. Sci Rep 8, 11531 (2018).

25. M. Lucanic, W. T. Plummer, J. Harke, M. Lucanic, E. Chen, D. Bhaumik, G. Harinath, A. Coleman-Hulbert, K. Dumas, B. Onken, E. Johnson, A. Foulger, S. Guo, A. Crist, M. Presley, J. Xue, C. Sedore, M. Chamoli, C. Chang, M. Chen, S. Angeli, M. A. Royal, J. Willis, D. Edgar, S. Patel, E. Chao, S. Kamat, J. Hope, C. Ibanez-Ventoso, J. Kish, M. Guo, P. Phillips, G. Lithgow, M. Driscoll, Standardized Protocols from the Caenorhabditis Intervention Testing Program 2013-2016: Conditions and Assays used for Quantifying the Development, Fertility and Lifespan of Hermaphroditic Caenorhabditis Strains. Protocol Exchange, doi: 10.1038/protex.2016.086 (2017).

26. S. Kim, S. Beak, S. Park, Supplementation with Triptolide Increases Resistance to Environmental Stressors and Lifespan in *C. elegans*. Journal of Food Science 82, 1484–1490 (2017).

27. L. J. Zhang, O. Elsallabi, C. Soto-Palma, J. Bartz, R. Salekeen, A. Nunes, W. Xu, K. Lee, B. Hughes, B. Zhang, A. Mohamed, S. J. McGowan, L. Angelini, R. O’Kelly, S. A. Biashad, E. Hillpot, F. Morandini, A. Seluanov, V. Gorbunova, X. Dong, L. J. Niedernhofer, P. D. Robbins, Fucoidans are senotherapeutics that enhance SIRT6-dependent DNA repair. Cell Biology [Preprint] (2025). 10.1101/2025.04.27.650852.

28. C. E. Burd, M. S. Gill, L. J. Niedernhofer, P. D. Robbins, S. N. Austad, N. Barzilai, J. L. Kirkland, Barriers to the Preclinical Development of Therapeutics that Target Aging Mechanisms: Table 1. GERONA 71, 1388–1394 (2016).

29. S. Angeli, I. Klang, R. Sivapatham, K. Mark, D. Zucker, D. Bhaumik, G. J. Lithgow, J. K. Andersen, A DNA synthesis inhibitor is protective against proteotoxic stressors via modulation of fertility pathways in Caenorhabditis elegans. Aging 5, 759–769 (2013).

30. K. R. Kasimatis, M. J. Moerdyk-Schauwecker, P. C. Phillips, Auxin-Mediated Sterility Induction System for Longevity and Mating Studies in. (2018).

31. K. Avchaciov, K. J. Clay, K. A. Denisov, O. Burmistrova, M. Petrascheck, P. O. Fedichev, AI -Driven Identification of Exceptionally Efficacious Polypharmacological Compounds That Extend the Lifespan of *Caenorhabditis elegans*. Aging Cell 24, e70060 (2025).

32. K. J. Clay, M. Sanchez-Alavez, I. Newman, N. Na, A. P. Verduzco Espinoza, A. To, S. Saad, H. T. Cline, M. Petrascheck, Atypical tetracyclines promote longevity and ferroptotic neuroprotection via translation attenuation. Pharmacology and Toxicology [Preprint] (2026). 10.64898/2026.01.09.698733.

33. J. H. Hartman, S. J. Widmayer, C. M. Bergemann, D. E. King, K. S. Morton, R. F. Romersi, L. E. Jameson, M. C. K. Leung, E. C. Andersen, S. Taubert, J. N. Meyer, Xenobiotic metabolism and transport in *Caenorhabditis elegans*. Journal of Toxicology and Environmental Health, Part B 24, 51–94 (2021).

34. M. Lucanic, W. T. Plummer, E. Chen, J. Harke, A. C. Foulger, B. Onken, A. L. Coleman-Hulbert, K. J. Dumas, S. Guo, E. Johnson, D. Bhaumik, J. Xue, A. B. Crist, M. P. Presley, G. Harinath, C. A. Sedore, M. Chamoli, S. Kamat, M. K. Chen, S. Angeli, C. Chang, J. H. Willis, D. Edgar, M. A. Royal, E. A. Chao, S. Patel, T. Garrett, C. Ibanez-Ventoso, J. Hope, J. L. Kish, M. Guo, G. J. Lithgow, M. Driscoll, P. C. Phillips, Impact of genetic background and experimental reproducibility on identifying chemical compounds with robust longevity effects. Nat Commun 8, 14256 (2017).

35. J. D. Rothstein, S. Patel, M. R. Regan, C. Haenggeli, Y. H. Huang, D. E. Bergles, L. Jin, M. D. Hoberg, S. Vidensky, D. S. Chung, S. V. Toan, L. I. Bruijn, Z. Su, P. Gupta, P. B. Fisher, b-Lactam antibiotics offer neuroprotection by increasing glutamate transporter expression. 433 (2005).

36. G. B. Phelps, J. Morin, C. Pinto, L. Schoenfeldt, S. Guilmot, A. Ocampo, K. Perez, Comprehensive evaluation of lifespan-extending molecules in C. elegans. [Preprint] (2024). 10.1101/2024.06.24.600458.

37. A. Griffin, K. R. Hamling, K. Knupp, S. Hong, L. P. Lee, S. C. Baraban, Clemizole and modulators of serotonin signalling suppress seizures in Dravet syndrome. Brain, aww342 (2017).

38. L. Zhang, L. E. Pitcher, V. Prahalad, L. J. Niedernhofer, P. D. Robbins, Targeting cellular senescence with senotherapeutics: senolytics and senomorphics. The FEBS Journal 290, 1362–1383 (2023).

39. A. Palazzo, G. Makulyte, D. Goerhig, J.-J. Médard, V. Gros, F. Trottein, S. Adnot, D. Vindrieux, J.-M. Flaman, D. Bernard, Benidipine calcium channel blocker promotes the death of cigarette smoke-induced senescent cells and improves lung emphysema. Aging 15, 13581–13592 (2023).

40. L. P. O’Reilly, C. J. Luke, D. H. Perlmutter, G. A. Silverman, S. C. Pak, C. elegans in high-throughput drug discovery. Advanced Drug Delivery Reviews 69–70, 247–253 (2014).

41. C.-L. Sun, H. Zhang, M. Liu, W. Wang, C. M. Crowder, A screen for protective drugs against delayed hypoxic injury. PLoS ONE 12, e0176061 (2017).

42. M. J. Smout, A. C. Kotze, J. S. McCarthy, A. Loukas, A Novel High Throughput Assay for Anthelmintic Drug Screening and Resistance Diagnosis by Real-Time Monitoring of Parasite Motility. PLoS Negl Trop Dis 4, e885 (2010).

43. M. Zamanian, J. D. Chan, High-content approaches to anthelmintic drug screening. Trends in Parasitology 37, 780–789 (2021).

44. H. Kadri, O. A. Lambourne, Y. Mehellou, Niclosamide, a Drug with Many (Re)purposes. ChemMedChem 13, 1088–1091 (2018).

45. G. McColl, B. R. Roberts, T. L. Pukala, V. B. Kenche, C. M. Roberts, C. D. Link, T. M. Ryan, C. L. Masters, K. J. Barnham, A. I. Bush, R. A. Cherny, Utility of an improved model of amyloid-beta (Aβ1-42) toxicity in Caenorhabditis elegans for drug screening for Alzheimer’s disease. Mol Neurodegeneration 7, 57 (2012).

46. S. Brenner, THE GENETICS OF *CAENORHABDITIS ELEGANS*. Genetics 77, 71–94 (1974).

47. M. Porta-de-la-Riva, L. Fontrodona, A. Villanueva, J. Cerón, Basic Caenorhabditis elegans Methods: Synchronization and Observation. JoVE, 4019 (2012).

48. S. Chilamkurthy, Transfer Learning for Computer Vision Tutorial (2017). https://docs.pytorch.org/tutorials/beginner/transfer_learning_tutorial.html.

